# Recognition of histone H3 methylation states by the PHD1 domain of histone demethylase KDM5A

**DOI:** 10.1101/715474

**Authors:** James E Longbotham, Mark J S Kelly, Danica Galonić Fujimori

## Abstract

PHD reader domains are chromatin binding modules often responsible for the recruitment of large protein complexes that contain histone modifying enzymes, chromatin remodelers and DNA repair machinery. A majority of PHD domains recognize N–terminal residues of histone H3 and are sensitive to the methylation state of Lys4 in histone H3 (H3K4). Histone demethylase KDM5A, an epigenetic eraser enzyme that contains three PHD domains, is often overexpressed in various cancers and its demethylation activity is allosterically enhanced when its PHD1 domain is bound to the H3 tail. The allosteric regulatory function of PHD1 expands roles of reader domains, suggesting unique features of this chromatin interacting module. Our previous studies determined the H3 binding site of PHD1, although it remains unclear how the H3 tail interacts with the N–terminal residues of PHD1 and how PHD1 discriminates against H3 tails with varying degrees of H3K4 methylation. Here we have determined the solution structure of apo and H3 bound PHD1. We observe conformational changes occurring in PHD1 in order to accommodate H3, which interestingly binds in a helical conformation. We also observe differential interactions of binding residues with differently methylated H3K4 peptides (me0, me1, me2 or me3), providing a rational for this PHD1 domain’s preference for lower methylation states of H3K4. We further assessed the contributions of various H3 interacting residues in the PHD1 domain to the binding of H3 peptides. The structural information of the H3 binding site could provide useful information to aid development of allosteric small molecule modulators of KDM5A.

## Introduction

Histone modifications play a central role in regulating chromatin structure and function. Protein reader domains recognize specific histone modifications and are often responsible for recruiting large protein complexes that contain histone modifying enzymes, chromatin remodelers and DNA repair machinery^1^. Plant homeodomain (PHD) domains are specific for recognizing the N–terminal residues of histone H3 and read the methylation state of H3K4 and, to a lesser extent, H3R2^2^. While the majority of PHD domains recognize H3, recognition of H4 peptides has been observed in extended PHD domains in MLL3/4^3–5^.

The conserved PHD fold consists of a two strand β–sheet, a C–terminal α–helix and two structural zinc atoms stabilized by a Cys4–His–Cys3 motif^2^. Depending on the residues involved, different PHD domains preferentially engage H3 as a function of the methylation status of H3K4. H3K4me3 preferring PHD domains, such as BPTF^6^ and TAF3^7^, have two to four aromatic residues that form an aromatic cage in the H3K4 binding pocket which allows for cation–π interactions with tri–methylated H3K4^8^. Inclusion of a H3K4 interacting glutamate residue in the aromatic cage of PHF20–PHD leads to the specificity for H3K4me2^9^. PHD domains that preferentially bind unmodified H3K4, such as BHC80^10^ and AIRE^11,12^, lack an aromatic cage, but instead have a H3K4 binding pocket consisting of both polar and hydrophobic residues. Specificity for unmodified H3K4 is achieved by electrostatic and hydrogen bond interactions. These domains discriminate against methylated H3K4 residues by steric occlusion of H3K4 methyl groups from the binding pocket in addition to an increase in cationic radius and decrease in hydrogen bond potential^10–15^.

Members of the KDM5 family are H3K4me3 specific demethylases and contain varying number of PHD domains. All four members of the KDM5 family are involved in pathogenesis and progression of various cancers^16–31^ and other, particularly neurological, disorders^32,33^. For this reason there is significant interest in developing therapeutics targeting these enzymes^34–38^. KDM5A specifically is overexpressed in breast cancer, where it promotes cancer progression and metastasis^39^, and there is evidence for its role in resistance to endocrine therapy^40^ as well as cancer drug resistance in lung cancer models^41^. KDM5A contains three PHD domains (**Figure 1a**). Its PHD3 domain is specific for H3K4me3 and it is thought to be responsible for recruiting KDM5A to the site of its substrate. PHD3 is also known to form a fusion with NUP98 in acute leukemia^16^. We have previously shown that the PHD1 domain preferentially binds unmodified H3^42^ and that the engagement of the PHD1 domain with its ligand allosterically enhances the demethylation activity of KDM5A by stabilizing substrate binding to the catalytic domain^43^. Despite the importance of PHD1 in the demethylase activity of KDM5A, there is limited structural information on how its PHD1 domain interacts with H3 and how it can discriminate against the methylation states of H3K4 peptides. We have previously mapped the unmodified H3 peptide binding site of PHD1 using NMR^42^. Important structural insights have also been gained from studies performed on the PHD1 domain of homologue KDM5B^13,14^. However, questions remain about the interactions of the N–terminal region of the PHD1 domain, which was absent in the construct used in our previous NMR studies, as well as how peptide binding residues of this reader domain interact with methylated H3 peptides.

**Figure 1.**
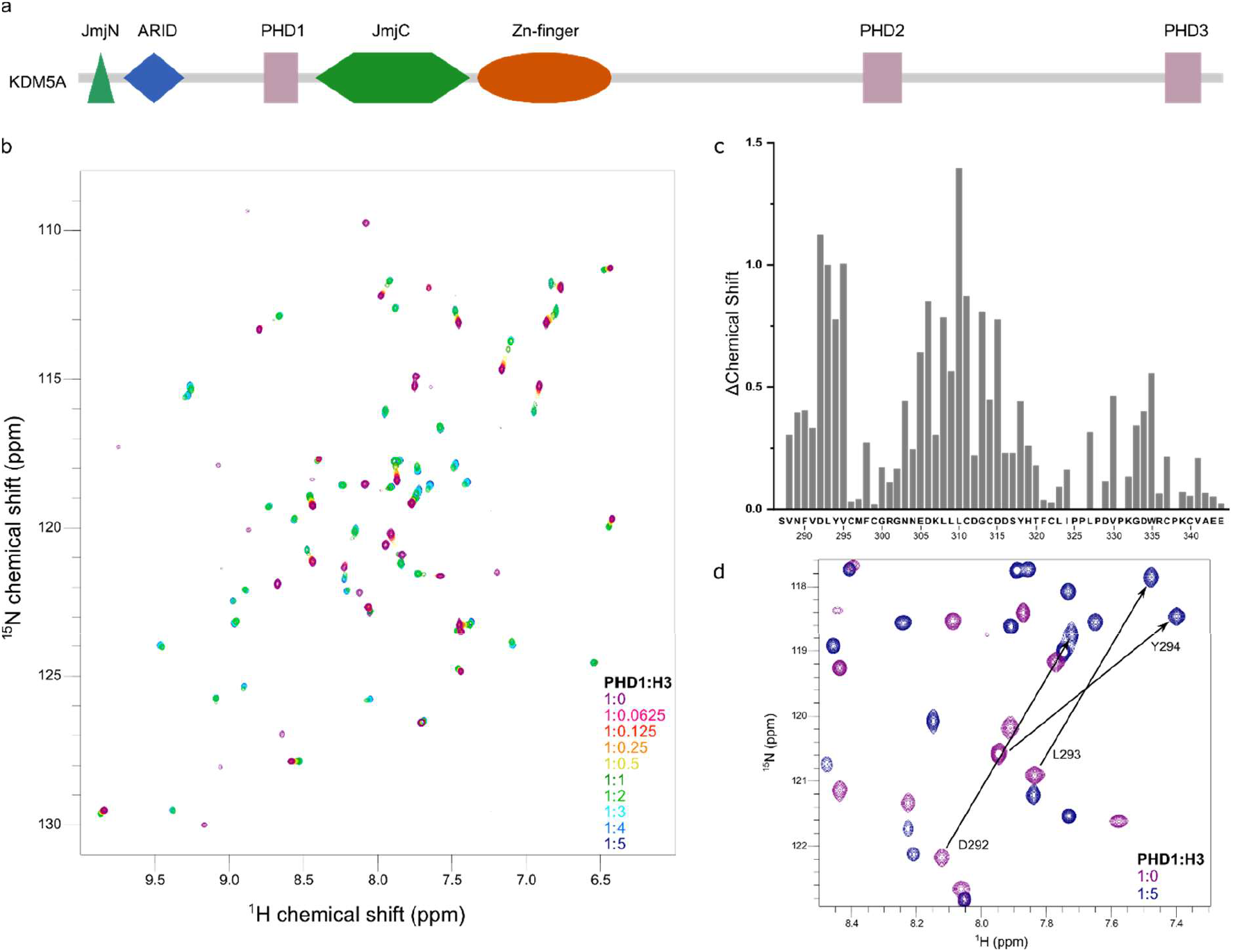
The N–terminal region of PHD1 undergoes large chemical shift perturbations upon H3 peptide binding. a) Domain architecture of KDM5A. b) 2D HSQC spectra of H3 10mer titrations with ^15^N labelled PHD1 (S287–E344). c) Chemical shift perturbations of peaks shown in b). d) Chemical shift perturbation of D292-Y294 upon addition of H3 10mer peptide. Apo PHD1 peaks are in magenta and H3 10mer bound peaks in blue.

Here we describe the NMR solution structure of PHD1 in apo and H3 bound forms. We observe that H3 adopts a helical conformation when bound to PHD1, allowing for additional H3 side chain interactions distal to those enabled by the critical H3 N-terminal residues. In addition, we find that H3K4 forms unique interactions with the N-terminal region of PHD1, not previously seen in the PHD1 domains of the KDM5 family. While unstructured in its apo form, the N-terminal region of PHD1 adopts a helical turn to make these interactions possible. The interactions between H3 peptide and PHD1 are modulated by the methylation status of H3K4, as we observe differential engagement of the N-terminal region of PHD1 with differently methylated H3 peptides. Peptide binding residues in PHD1 were further probed through mutagenesis. Structural information on ligand engagement by this reader domain has revealed a role of secondary structure adopted by histone ligand to facilitate additional interactions. Additionally, given the allosteric role of PHD1 in regulation of KDM5A catalysis, structural information could assist in the development of unique allosteric small molecule ligands for use as chemical tools. This would allow for pharmacological interrogation of the role of this enzyme in KDM5A dependent disorders.

## Methods

### Expression and purification of WT and mutant GST–PHD1

WT and mutant PHD1 constructs (S287–K347) were cloned into a pET41a vector and expressed in BL21(DE3) *E. coli* cells. Expression and purification of WT and variants followed the same protocol. Cells were induced with 0.4 mM IPTG and grown at 18 °C overnight before the pellet was collected. The cells were resuspended in lysis buffer (140 mM NaCl, 2.7 mM KCl, 10 mM Na_2_HPO_4_, 1.8 mM KH_2_PO_4_, 5 mM β–mercaptoethanol, 50 μM ZnCl_2_, 1 mM phenylmethylsulfonyl fluoride (PMSF), pH 7.3), lysed by sonication and centrifuged. The supernatant was purified using Glutathione Sepharose 4B resin, washed with high salt buffer (50 mM HEPES, 700 mM KCl, 10% glycerol, 5 mM β–mercaptoethanol, 50 μM ZnCl_2_, 1 mM PMSF, pH 8.0) and low salt buffer (50 mM HEPES, 150 mM KCl, 10% glycerol, 5 mM β– mercaptoethanol, 50 μM ZnCl_2_, 1 mM PMSF, pH 8) and recovered by elution using low salt buffer supplemented with 30 mM glutathione. The sample was then further purified by size– exclusion chromatography using a Hiload 26/60 Superdex 75 gel filtration column in a buffer of 40 mM HEPES, 50 mM KCl, 5 mM β–mercaptoethanol, and 50 μM ZnCl_2_, pH 8.0 concentrated and flash–frozen. 1D ^1^H NMR was used to conform the correct folding of mutants (**Supplementary Figure 1**)

### Fluorescence polarization binding assays

The binding of GST–PHD1 to H3 10mer peptides was measured by either direct or competition–based fluorescence polarization (FP). For direct FP binding assay, 10 nM of C– terminal fluorescein labeled H3 10mer peptide were incubated for 30 mins with varying concentrations of GST–PHD1. Data from direct FP measurements were fitted to equation 1:

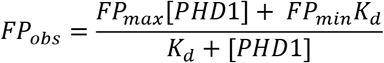

For competition–based FP assays, GST–PHD1, at a concentration equal to the *K*_d_ for unmodified H3 peptide, was incubated with 10 nM of C–terminal fluorescently labeled H3 10mer peptide and different concentrations of unlabeled peptides were used as competitors. Data from competition–based FP assays were fitted to equation 2:

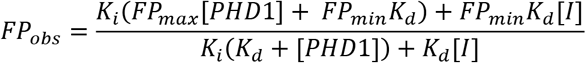

where FP_obs_ is the observed FP, FP_max_ is the maximum FP value, FP_min_ is the minimum FP value, [PHD1] is concentration of GST–PHD1, *K*_d_ is the dissociation constant, *K*_i_ is the inhibition constant referring to the competing peptide and [I] is the competing peptide concentration.

### NMR binding experiments

A PHD1 construct encompassing residues S287–E344 was used in NMR studies. The GST– PHD1 fusion construct was expressed in M9 minimal media containing ^15^N ammonium chloride and/or ^13^C_6_ D–glucose. The labelled GST–PHD1 was purified as described for binding studies with additional steps of removing the GST tag by TEV protease cleavage after elution from the Glutathione 4b resin. After cleavage, the sample was incubated with nickel resin to remove the His-tagged TEV and the flow–through, containing the cleaved PHD1, was collected. Finally, the cleaved PHD1 was loaded onto a Hiload 26/60 Superdex 75 gel filtration column pre–equilibrated with 50 mM HEPES, 150 mM NaCl, 0.5 mM TCEP and 0.1 mM ZnCl_2_ at pH 7.5.

550 μM ^13^C-, ^15^N-labelled PHD1 was used in 3D triple–resonance experiments for the backbone assignment of apo–PHD1. For the assignment of H3 10mer bound PHD1, 800 μM H3 10mer was added to the sample. 3D CBCANH and CBCA(co)NH spectra were recorded on a Bruker 500 MHz AVANCE DRX spectrometer equipped with an actively shielded Z– gradient QCI cryoprobe (^15^N/^13^C/^31^P, ^1^H) using programs from the pulse program library (TopSpin 1.3pl10).The 3D spectra were processed in NMRPipe and proton chemical shifts were referenced externally to a DSS standard and while ^13^C and ^15^N chemical shifts were referenced indirectly to this value^44,45^.

For 2D ^15^N–HSQC peptide titration experiments, 200 μM ^15^N–labeled PHD1 was used and spectra measured after each addition of H3 10mer peptides. The 2D ^15^N-FHSQC spectra for the H3 10mer peptide titration series were recorded on a Bruker 800 MHz AVANCE-I spectrometer equipped with an actively shielded Z-gradient TXI cryoprobe using programs from the pulse program library (TopSpin 2.1). Chemical shift perturbations of HSQC peaks, upon addition of H3 10mer peptide, were calculated with the following equation:

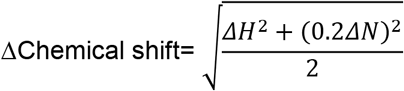

All experiments were carried out at 298 K (calibrated with 4% v/v MeOH in MeOD; using the following coefficients of T (K) = (4.109 - Δ) * 0.008708 where Δ is the chemical shift difference between the CH_3_ and OH protons in methanol) in 50 mM HEPES, 150 mM NaCl, 0.5 mM TCEP and 0.1 mM ZnCl_2_ at pH 7.5 in 5% D_2_O.

### NMR structure determination

In order to determine the solution structure of apo PHD1 900 μM ^13^C-, ^15^N-labelled PHD1 (S287-E344) was used. For the H3 bound structure 900 μM ^13^C-,^15^N-labelled PHD1 (S287-E344) was used, but with addition of 4 mM H3 (1-10). Spectra were recorded on a 600 MHz Bruker Avance NEO spectrometer equipped with an actively shielded Z-gradient 5mm TCI cryoprobe using programs from the pulse program library (TopSpin 4.0.6). The backbone assignments for free and bound PHD1 were extended to the sidechain resonances using 2D ^15^N-FHSQC, 2D ^13^C-HSQC, 2D ^13^C-CTHSQC, 3D H(ccco)HN-TOCSY, 3D h(Ccco)NH-TOCSY and 3D HCcH-TOCSY experiments. Spectra were processed in TopSpin 4.0.7 and analyzed using CcpNmr Analysis 2.4^46^. For the Apo PHD1 domain distance restraints were measured using a 120 ms simultaneous evolution 3D ^13^C/^15^N-NOESY-HSQC experiment^47^. For the H3-bound PHD1, distance restraints were measured using a 3D ω1-^13^C/^15^N-filtered, ω3 ^13^C-/^15^N-edited NOESY-HSQC experiment with simultaneous evolution^48^. The spectrum was split by adding or subtracting components to generate two sub-spectra containing intramolecular (3D ω3-^13^C/^15^N-edited NOESY-HSQC) and intermolecular nOes (3D ω1-^13^C/^15^N-filtered, ω3-^13^C/^15^N-edited NOESY-HSQC). To assign the resonances of the H3 peptide bound to PHD1 40 ms ω1, ω2-^13^C/^15^N-doubly filtered TOCSY and 120 ms ω1, ω2-^13^C/^15^N-doubly filtered NOESY experiments were recorded^45^. Whereas a sample containing 900 μM ^13^C-,^15^N-labelled PHD1 (S287-E344) with 4 mM H3 was used in the 3D filtered/edited NOESY experiments to obtain intermolecular distance restrains, in the 2D doubly filtered NOESY intramolecular nOes signals from the bound peptide were strongest with a lower peptide to protein ratio (1250 μM ^13^C-,^15^N-labelled PHD1 and 400 μM H3).

Dihedral restraints were generated using the program DANGLE^49^. Hydrogen bond restraints included in the structure calculations were based on measurements of amide chemical exchange with solvent detected by 2D ^15^N-CLEANEX-HSQC experiments (using mixing times of 5, 20 and 100 ms)^45^, secondary structure predicted by DANGLE and nOe patterns typical of α-helical secondary structure.

The programs ARIA version 2.3.2^50^ and CNS version 1.2.1^51^ were used to calculate the NMR structures. To solve the structures, 9 iterations of simulated annealing were performed using CNS. For the first 8 rounds of simulated annealing, the n_structures parameter was set to 50 and the n_best_structures parameter was set to 15. For the 9^th^ round, the n_structures parameter was set to 100 and the n_best_structures parameter was set to 15. Finally, a refinement in water was performed on the 25 lowest energy structures from the 9^th^ iteration. Initial structure calculations were conducted without hydrogen bond restraints. Hydrogen bond donors were then identified, and the corresponding hydrogen bond restraints included in later calculations. Calculations were initially conducted using a flat-bottom harmonic wall energy potential for the distance restraints. The final calculation was then performed using a log-harmonic potential^52^ with Bayesian weighting of the restraints^53^ and a softened force-field^54^. Structures were validated using the Protein Structure Validation Software (PSVS) suite 1.5^55^. The chemical shifts, restraints and structural co-ordinates have been deposited with the BMRB (Apo - 30808; H3 bound - 30809) and PDB (Apo - 7KLO, H3 bound - 7KLR).

## Results and Discussion

### Unstructured N–terminal residues participate in H3 peptide recognition

PHD1 preferentially binds unmodified H3 peptides and binding affinity decreases with an increase in H3K4 methylation^42,43^. We previously mapped the H3 peptide binding site of PHD1_KDM5A_ domain using two–dimensional heteronuclear single quantum coherence (2D ^15^N– HSQC) NMR titration experiments^42^. However, these experiments did not detect any interaction between the H3 peptide and D292, a H3K4 binding pocket residue observed in close homologue KDM5B, in the PHD1 construct used (D292–E344). Our recent fluorescence polarization–based binding studies suggested that D292 is involved in binding when an N– terminally extended PHD1 construct (S287–K347) is used^43^. This observation prompted us to examine the contribution of these N–terminal residues to peptide binding and, additionally, to H3K4 methyl recognition.

To assess binding of the H3 peptide to PHD1, 2D ^15^N HSQC experiments were conducted using a PHD1, S287–E344, construct. The HSQC spectrum of apo PHD1 showed a well– dispersed set of peaks, consistent with a well–structured domain, which were assigned to individual residues using NMR triple–resonance backbone experiments (**Figure 1b**). Comparison of the HSQC spectra of S287–E344 with D292–E344, which we previously used, showed several overlapping peaks, but a significant shift in many residues suggests structural differences between the two constructs (**Supplementary Figure 2a**). These differing peaks include L293-V295, G302-E305, L309-D316, I324, D334, W335, V341. When mapped onto the structure of apo PHD1 these residues cluster in the N-terminal and structurally adjacent regions of the domain (**Supplementary Figure 2b**). These observations suggest that the presence of five additional N-terminal residues impacts the structure of the domain beyond its N-terminus.

The H3 10mer peptide binding site in the S287–E344 PHD1 construct was identified by NMR peptide titration experiments (**Figure 1b,c**). Large chemical shift changes were observed on binding, many undergoing slow–intermediate exchange on the NMR timescale, indicating a strong interaction (low–micromolar affinity). The largest shifts were found to occur in the N– terminal region (D292–V295), residues preceding the first β–strand (E305, D306), the first β– strand itself (L308–C311) and following residues, which are part of the second Zn finger (G313, D315) (**Figure 1c**). Additionally, residues near the C–terminal were also perturbed (V330, W335). Large shifts in D292-V295, and their adjacent residues, not seen in studies with D292–E344 PHD1, indicate the importance of these residues in H3 binding. Similar trends have been observed in NMR studies of AIRE^11^ and the PHD1_KDM5B_^13,14^.

In the absence of the H3 ligand, the chemical shifts values of the N–terminal residues (V288– Y294) of PHD1 suggest that they are unstructured. However, upon addition of H3 10mer peptide we observe shifts in D292–Y294 away from unstructured chemical shift values, likely due to structural reorganization (**Figure 1d**). Secondary structure prediction suggested the formation of a helix in the N-terminal region (N289-D292) when H3 peptide is bound (**Supplementary Figure 3**). Ligand–induced structural changes have been previously observed in the PHD domain of PHF20^9^. To gain further insight into the structure of PHD1_KDM5A_ and interactions with H3, we determined the NMR structures of apo and H3 bound PHD1.

### H3 adopts a helical conformation upon binding to PHD1

We determined the solution structures of apo PHD1 and PHD1 bound to a H3 10mer (A1-S10) using triple resonance NMR methods (**Supplementary Table 1**). Both the apo and H3 bound structures adopt the conserved PHD fold where residues L308-L310 and S317-H319 form an anti-parallel β-sheet and residues P338-E343 form a C-terminal α-helix (**Figure 2a, b**). H3 binds to an acidic groove and unexpectedly adopts an α-helical conformation (**Figure 2b, c**). The N-terminal residues of the H3 peptide, A1-K4, bind to PHD1 in a similar manner to other PHD domains^2^. The NH_2_-terminal amine group of H3 interacts with the carbonyl groups of P331 and G333 and the side chain of H3A1 sits in a hydrophobic pocket formed by L309, P331 and W335 (**Figure 2d**). In addition, the H3R2 carbonyl hydrogen bonds with the amide of L310 and the amide of H3K4 forms hydrogen bonds with the carbonyl of L308 (**Figure 2d, e**). The H3R2 side chain interacts with D312, a conserved residue we have previously shown is critical for affinity^42^, and forms hydrophobic interactions with L310. In addition, D315 is shown to contribute to the acidic R2 binding pocket, but it is part of a flexible loop which orientates towards R2 in a subset of the ensemble structures. This is in agreement with diminished H3 binding observed in a previously studied D315A mutant^42^. The H3K4 side chain extends from the base of the H3 helix into the acidic groove. The K4 side chain forms charge-charge interactions with the side chain of E305 and hydrogen bonds with the carbonyl groups of V291, D292 and Y294. Hydrophobic interactions are also observed with the side chain of L308. (**Figure 2f**). Interaction with these mainchain atoms of V291, D292 and Y294 in the N-terminal region stabilizes a helical turn conformation in V291-L293. This agrees with our secondary structure prediction data (**Supplementary Figure 3**) which suggested formation of a helix in the N-terminal region of PHD1 upon H3 binding.

**Figure 2.**
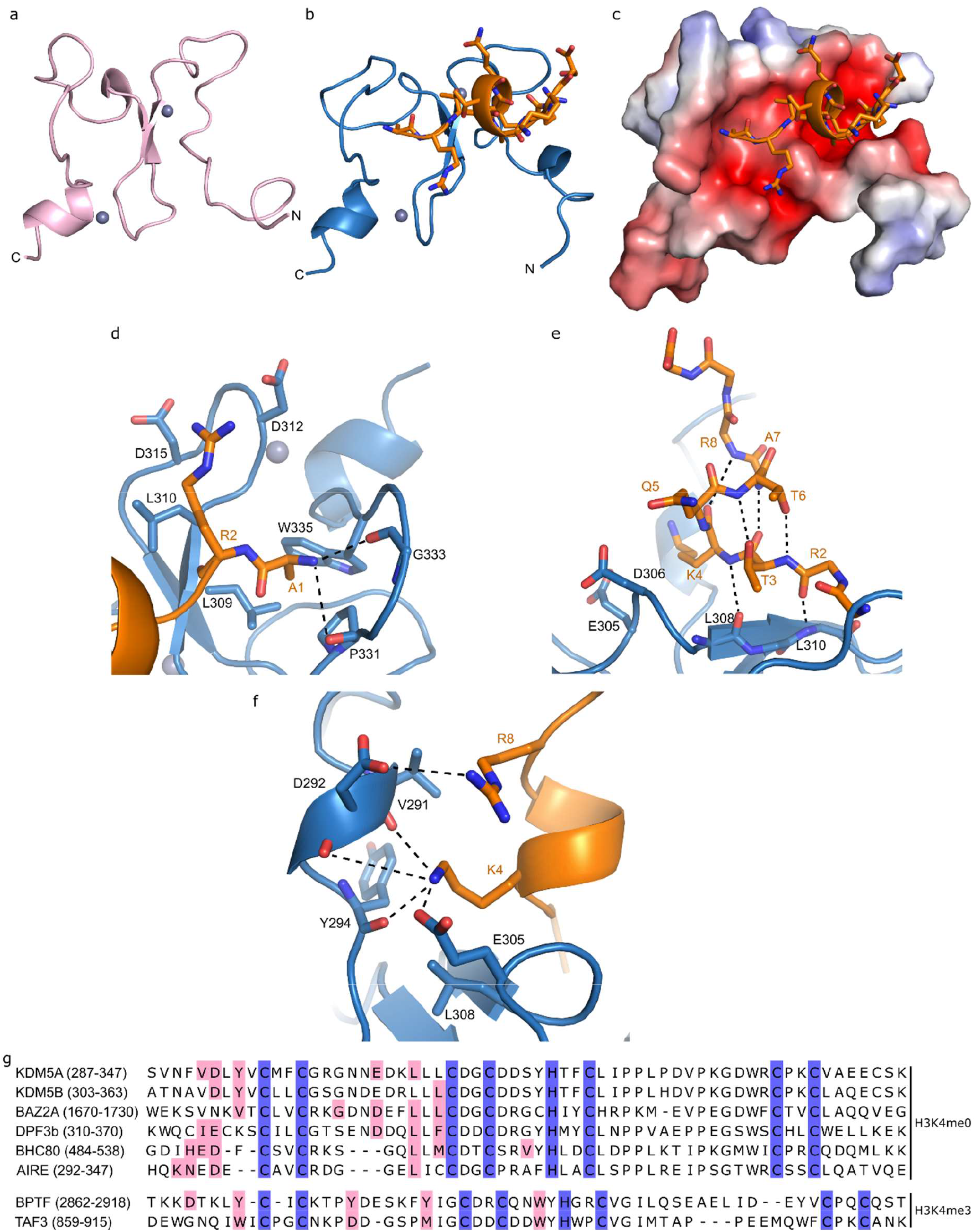
Structure of the PHD1 domain of histone demethylase KDM5A. a) Apo structure of PHD1 (PDB: 7KLO), shown in pink. b) H3 10mer bound structure of PHD1 (PDB: 7KLR). PHD1 is shown in blue, H3 is shown in orange. c) Electrostatic potential surface of PHD1 calculated with APBS. Red indicates a negative potential and blue indicates positive potential. d) H3A1 and H3R2 binding pocket. e) Helical conformation and backbone interactions of H3 with PHD1. A1, R2, A7-S10 side chains are omitted for clarity. f) H3K4 binding pocket. g) Sequence alignment of H3K4me0 and H3K4me3 preferring PHD1 domains. Residues highlighted in pink interact with H3K4. Conserved Zinc finger residues are shown in blue.

Residues H3K4-R8 adopt a helical conformation, which is stabilized by intramolecular H3 hydrogen bonds between the carbonyl groups of T3 and K4 to the amide protons of A7 and R8. We also observe hydrogen bonds from the side chain hydroxyl group of T3 to the amide of T6 and reciprocal hydrogen bonds from the T6 side chain hydroxyl group to the amide of T3 (**Figure 2e**). In addition, we observe intermolecular interactions that may stabilize the helical formation. The side chain of H3Q5 appears to form long hydrogen bonds with D306 and H3A7 and H3R8 form hydrophobic interactions with V291. As the H3 peptide turns, H3R8 is positioned above the H3K4 binding pocket where it forms charge interactions with the side chain of D292 (**Figure 2f**). We have previously shown that the affinity of H3R8me2s and H3R8me2a is reduced by two-fold compared to unmodified H3^43^, which suggests that methylation of H3R8 may impact this interaction and the helical conformation.

Several regions of PHD1 undergo conformational changes upon H3 binding (**Supplementary Figure 4**). These include the R2 binding pocket (C311-S317), where D312 and D315 orientate to interact with H3R2, and the initially unstructured residues in the K4 binding pocket (F290-V295). Interestingly, we also observe changes in a proline rich loop (C322-P328), distal to the H3 binding pocket. As PHD1 is a small domain it is likely that interaction of the H3R2 and H3K4 pockets with H3 pulls on the structural Zn fingers affecting regions distal to the H3 binding pocket. Given the role of PHD1 in allosteric regulation of KDM5A demethylase activity, it is plausible that these conformational changes might impact adjacent domains in the context of full length KDM5A and contribute to allosteric communication within the protein.

Overall, the structure of H3 bound PHD1 retains the conserved structural elements of PHD domains. However, the H3K4 binding pocket differs from that observed in the PHD1 domain of close homologue KDM5B, which shows interaction of the H3K4 with the side chains of D308, Y310 and L326^14^ (equivalent residues in KDM5A are D292, Y294 and L310). In KDM5A’s PHD1 structure we observe mainchain interactions with V291, D292, Y294 and side chain interactions with E305 and L308 (**Figure 2f**). H3K4 binding pockets in other PHD domains like AIRE^12^, BAZ2A^56^, DPF3b^57^ and BHC80^10^ have a more similar binding pockets to PHD1_KDM5A_, which consist of interactions with mainchain atoms and side chain interactions to equivalent residues to E305 and L308 (**Figure 2g**, **Supplementary Figure 5**).

H3 tail adopts helical conformation when bound to KDM5A’s PHD1 domain (**Figure 2**). Helical H3 conformations have been previously observed in the PHD domains of BAZ2A^56^, the TTD-PHD of UHRF1^58^, and the double PHD finger domains of DPF3b^59^, MOZ^60^ and MORF^61^. These domains bind H3 in a similar helical conformation to that observed in the structure of H3-bound PHD1_KDM5A_, including H3R8 interactions with acidic residues (**Supplementary Figure 6a-f**). In BAZ2A it has been shown that helicity is essential for H3 binding as it stabilizes the H3 peptide. In the case of PHD1_KDM5A_, the helical conformation allows for additional interactions between the ligand and reader domain, like those observed with H3R8. Bioinformatic analysis suggest that the presence of an acidic patch (residues corresponding to E305 and D306 in KDM5A) promotes the formation of a helical H3 by stabilizing the positive dipole of the N-terminus of H3^56^. This acidic patch is present in BAZ2A, UHRF1, DPF3b, MOZ and MORF as well as in KDM5B (**Figure 2g**). However, despite close sequence similarity between the PHD1 domains of KDM5A and KDM5B, PHD1_KDM5B_ binds to H3 in an extended conformation^14^ (**Supplementary Figure 6g**), a difference that may be attributed to difference in constructs used. In analogy, UHRF1 has been shown to bind H3 in helical and extended conformations. Helical H3 binding in UHRF1 was shown to require the TTD-PHD domain rather than isolated domains^58^.

Formation of a H3 helix is essential in bivalent recognition of histone modifications for certain reader domains. For example, the double PHD finger domains of DPF3b, MOZ and MORF require the helical H3 in order to orient the ligand to enable H3K14ac interactions^59–61^. In the TTD-PHD domain pair of UHRF1, the helical H3 conformation is necessary for interaction with H3K9me3^58^. H3 helicity has been observed up to H3T11, before a flexible G12, G13 patch in H3 binding to DPF3b, MOZ and MORF. We speculate that binding of H3 to PHD1 in a helical conformation in full length KDM5A could allow interaction of downstream H3 modifications with other KDM5A domains. It is interesting to note that the helical turn region (V291-L293) of PHD1 that interacts with the H3K4 has structural similarity to a turn in MOZ, MORF and DPF3b (**Supplementary Figure 6h**). In these double PHD finger domains this turn links to the additional PHD domain, which interacts with H3K14ac residues. While PHD1 is not a double PHD finger domain it is interesting to speculate whether ARID-PHD1 linker region residues N-terminal of the PHD1 construct used in this study could provide additional H3 interactions, like those observed in the double PHD finger domains.

### Differential engagement with methylated H3K4 peptides by PHD1

Methylation of H3K4 greatly impacts the affinity of the H3 peptide for PHD1 with an increase in the number of methyl groups on H3K4 resulting in a decreased affinity^42^. Interestingly, PHD1_KDM5A_ domain is still able to bind all three methylated states of H3K4, a feature distinct from several other PHD domains that engage unmodified H3 tails. For example AIRE^11^, TRIM24^62^, CHD4’s PHD2^63^, BHC80^10^ and the PHD–like domain of DNMT3L^64^ also show weak binding for methylated H3K4 peptides with *K*_d_ values >200 μM or non–binding for higher methylation states (H3K4me2 and me3). Even the closely related PHD1_KDM5B_ shows weaker binding (>80 μM for H3K4me2 and me3) to methylated H3K4 peptides^13,14^. The relatively high affinity of PHD1_KDM5A_ for H3K4 methylated peptides could explain how allosteric stimulation of KDM5A’s H3K4me3 demethylase activity initially occurs in H3K4me3 rich regions^43^. The ability of PHD1 to bind all three methylation states of H3K4 peptides, provides a unique opportunity to investigate how methylation is accommodated by this reader domain.

HSQC NMR titration measurements were performed with H3K4me1, H3K4me2 and H3K4me3 peptides (**Figure 3a-c**, **Supplementary Figure 7**). For all three methylated peptides we see the presence of NMR signals indicating fast exchange on the NMR timescale, suggesting weaker interactions than with unmodified H3. However, H3K4me1 still contains a large number of signals in slow– intermediate exchange on the NMR timescale suggesting strong interactions, comparable to with unmodified H3 (**Supplementary Figure 8**). We observe that similar residues are engaged upon binding of unmodified and methylated H3 peptides, implying an overall similar binding site (**Figure 3d**). However, significant differences in the behavior of specific residues show alternate interactions are likely made with methylated peptides.

**Figure 3.**
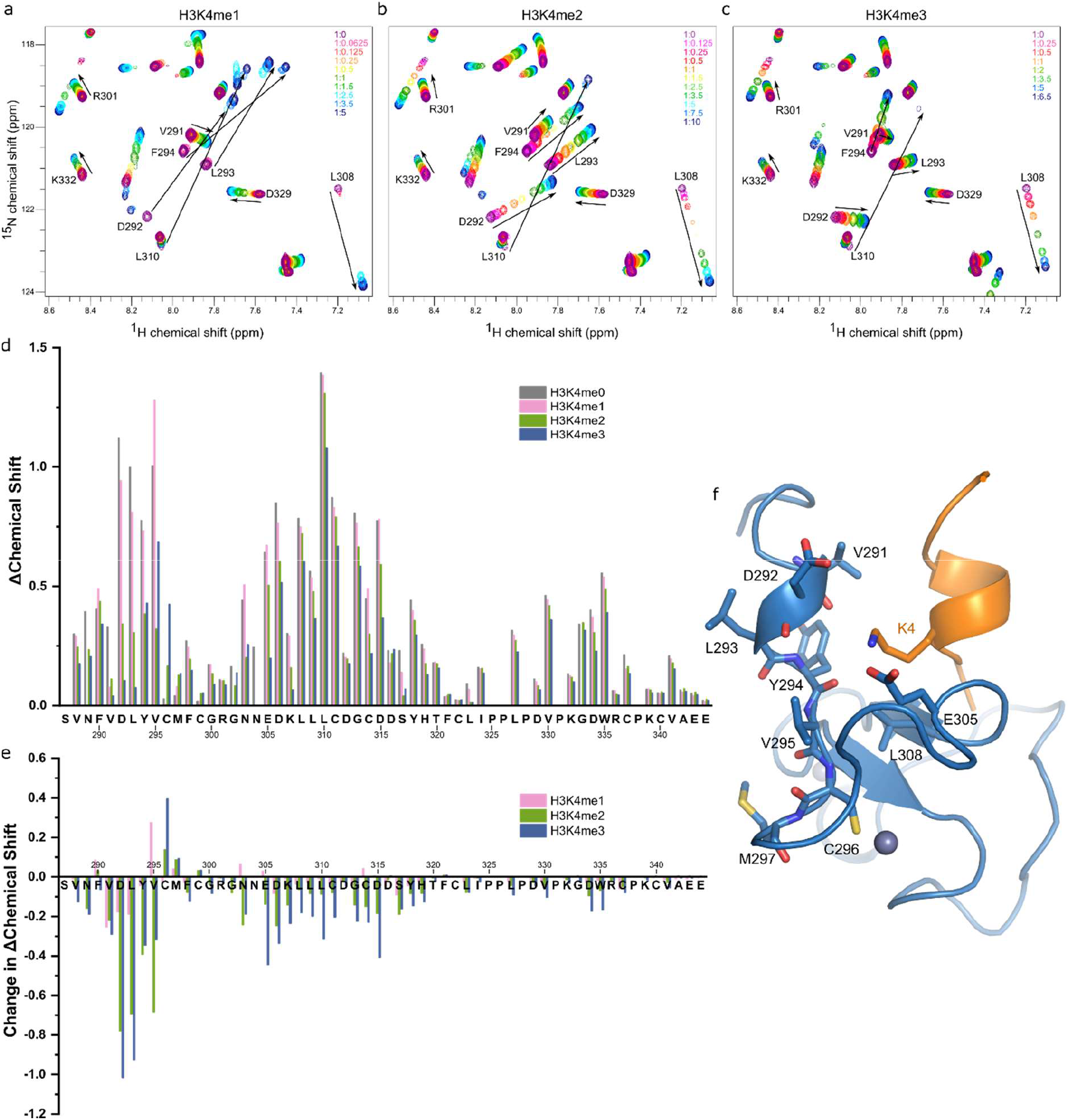
Binding to differently methylated H3 peptides causes reduced interaction with N– terminal residues. a–c) HSQC spectra of H3 10mer peptide titrations with ^15^N labelled PHD1 (S287– E344) and a) H3K4me1, b) H3K4me2 and c) H3K4me3. d) Chemical shift perturbation of peaks upon addition of H3K4me1, me2 and me3 10mer peptides. e) Change in chemical shift perturbations shown in d) relative to shifts in H3K4me0. f) H3K4 binding pocket and adjacent residues in PHD1.

When the H3K4me1 peptide is bound, the majority of peak shifts are similar to those on binding to the unmodified peptide (**Figure 3d, e; pink bars**). However, with mono–methylation some residues are affected to a lesser degree with the largest decreases in chemical shift changes occurring for residues V291–L293, residues part of helical turn that interacts with unmodified H3K4 (**Figure 3f**). Interestingly, other H3K4 binding pocket residues (Y294, E305 and L308) show relatively little change in chemical shift perturbations suggesting that mono–methylation does not affect their interaction with the H3K4 residue.

Smaller chemical shift changes, relative to those observed upon binding of unmodified H3 peptide, are observed with binding of the H3K4me2 peptide. The largest reduction in chemical shifts, relative to the unmodified H3 peptide, are in the V291–V295 region (**Figure 3d,e green bars**). Our observations suggest that the presence of two methyl groups causes reduced interactions with the N-terminal residues that interact with H3K4 (**Figure 3f**). These changes are in contrast to relatively small changes in residues E305 and L308, which appears to be unaffected by H3K4 di–methylation.

Binding of H3K4me3 causes a large reduction in the chemical shift perturbations, relative to unmodified H3. The largest differences in chemical shift changes between the unmodified and tri–methylated ligand occur in residues that form the H3R2 and H3K4 binding pockets (V291– V295 and E305–D315) (**Figure 3d, e; blue bars**). The reduced interaction of the V291–V295 region with the peptide is likely due to the bulky nature of the H3K4me3 side chain in the H3K4 binding pocket. Its presence is likely to prevent formation of interaction with the carbonyl groups of V291, D292 and Y294 and hinder optimal orientation of the helical turn. In addition, the increase in cationic radius and decrease in hydrogen bond potential would mean a reduced ability to interact with E305. Interestingly, upon H3K4me3 binding, C296 and M297 undergo increased perturbation compared to unmodified H3 binding. These residues are part of the first Zn–finger positioned a few residues down from the induced helical region (**Figure 3f**). We hypothesize that the N-terminal region is unstructured upon H3K4me3 binding and this flexibility may then put strain on the Zn finger stabilized C296 and M297.

Together our NMR data indicates that, of the H3K4 binding pocket residues, the N-terminal patch of H3K4 interacting residues (V291-Y294) are most sensitive to the methylation state of the peptides.

### Assessing contributions of H3 interacting residues

Our structural data have allowed us to understand the interactions of PHD1 and differently methylated H3 peptides. To further assess the contribution of the key H3 interacting residues in H3 peptide binding and methyl recognition we mutated these residues and measured H3 peptide binding to PHD1. Additionally, we analyzed known cancer-associated mutations in PHD1 to understand their impact on H3 affinity and methylation preference.

Mutation of D292 to alanine decreases the affinity of PHD1 for H3 ∼6–fold relative to WT (**Figure 4a**). Loss of interaction between D292 and H3R8 may disrupt the helical conformation of both H3 and V291-L293 and prevent optimal orientation of the H3K4 interacting carbonyls of V291, D292 and Y294 (**Figure 2f**). It also reduces binding to H3K4me1/2/3 peptides, with a 4–fold reduction for the interaction with H3K4me1 and ∼2–fold reduction for higher methylation states. This decreased dependence on the presence of D292 for binding higher methylation states of the peptide is consistent with our chemical shift perturbation measurements where we see a decreased interaction of D292 with methylated H3K4 (**Figure 3**).

**Figure 4.**
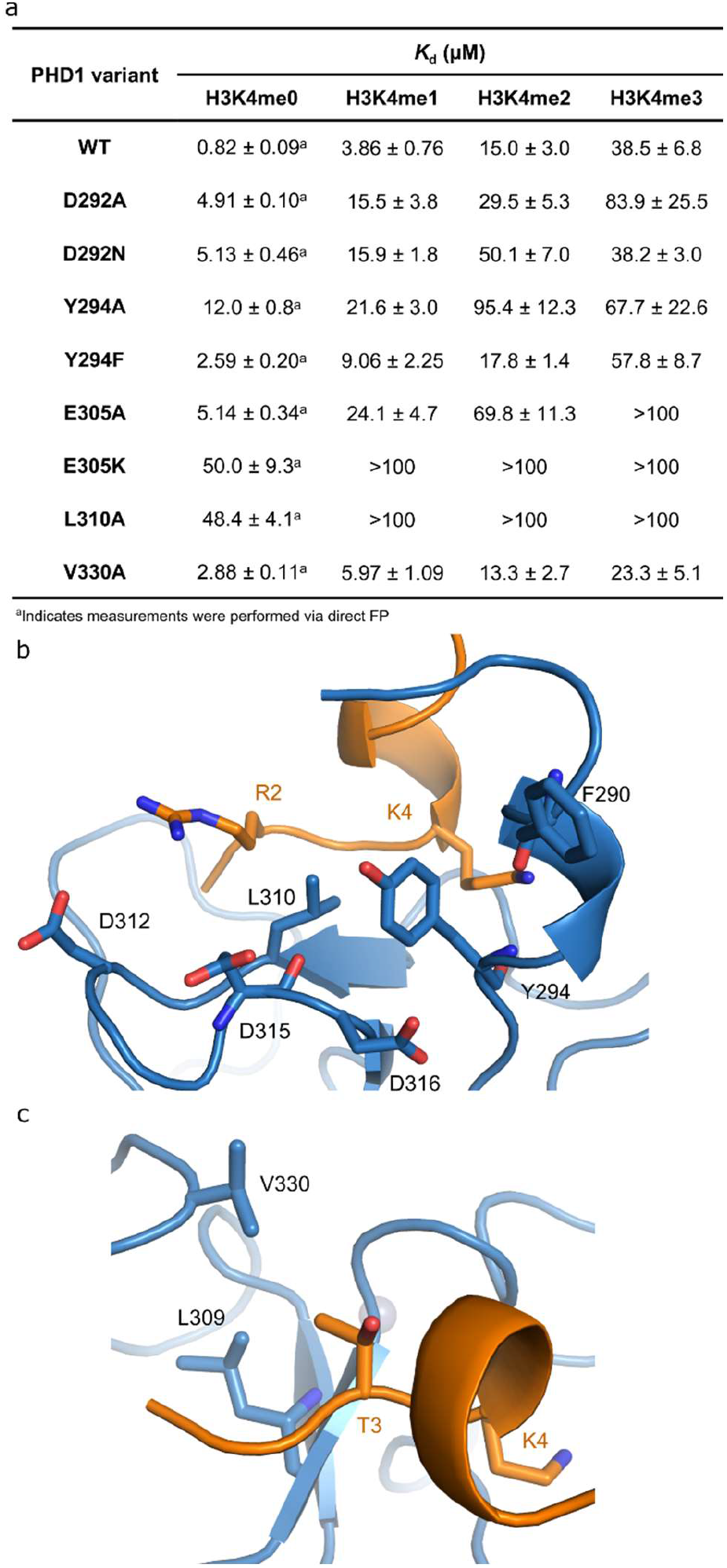
Binding affinities of PHD1 peptide binding site mutants for methylated H3 peptides. a) *K*_d_ values of PHD1 peptide binding pocket variants binding to differently methylated H3K4 10mer peptides. b) Y294 region of PHD1’s ligand binding pocket. c) H3T3 binding pocket.

Similar to D292A, cancer mutant D292N PHD1 has comparable fold–reductions in affinity for the unmodified and H3K4me1 peptides (**Figure 4a**). This is likely due to disruption of the helical turn region (V291-L293) that interacts with H3K4 (**Figure 2f**). *K*_d_ values for binding of the H3K4me2 peptide to the D292N mutant differ from those of D292A, but overall show a much reduced affinity relative to the WT PHD1. However, the *K*_d_ for H3K4me3 binding to the D292N mutant is similar to WT values, suggesting that the negative charge of the aspartate is not as important in H3K4me3 binding, but that other side chain contacts likely are.

E305 forms charged interactions with the H3K4 (**Figure 2f**). Mutation of this residue to Ala results in ∼6–fold reduction in affinity for both unmodified and mono–methylated H3K4 (**Figure 4a**) highlighting a significant contribution to affinity. The smaller decrease in affinity observed with H3K4me2 and me3 suggests a decreased dependence on interactions of these peptides with E305. These binding affinities agree with the reduced chemical shift perturbations of E305 when binding to H3K4me2 and me3 peptides (**Figure 3**). Cancer mutant E305K reverses the charge of this residue and perhaps expectedly we see large reduction in affinity, likely caused by electrostatic repulsion with H3K4 (**Figure 2f**).

L310A shows greatly reduced binding affinity for the unmodified and methylated H3 peptides, with a nearly 60–fold decrease in affinity for the unmodified H3 peptide. This is consistent with observations of a large chemical shift perturbation of L310 upon peptide binding. L310 interacts with H3R2 in the H3 peptide, which is a critical residue for peptide binding (**Figure 4b**).

V330 forms hydrophobic interactions with H3T3 and mutating this residue to an alanine reduced the affinity 3.5–fold for unmodified H3 (**Figure 4a, c**). Similar *K*_d_ values of the WT and V330A PHD1 towards methylated H3 peptides suggest that the interaction between V330 and H3T3 has a significant contribution only to the binding of unmodified H3.

We have previously shown that mutation of the conserved R2 interacting residues Asp312 and Asp315 greatly impacts binding, with Asp312 being essential for binding^42^. Additionally, we have shown that mutating of a conserved tryptophan residue W335 to alanine also greatly impacts binding as our structure shows that this is part of a hydrophobic pocket that interacts with H3A1 (**Figure 2d**).

Our investigation of various PHD1 cancer mutations could suggest their role in aberrant KDM5A function. Y294F is found in a bladder carcinoma; D292N in stomach adenocarcinoma and CNS glioma; E305K in breast invasive ductal carcinoma and head and neck squamous cell carcinoma; and V330M in cutaneous melanoma (COSMIC and cBioPortal databases^65–67^). We observed reduced binding capacity for low methylation–state H3K4 peptides with D292N and Y294F mutant PHD1 and show that E305K has the most striking impact on binding of H3 ligands in all methylation states (**Figure 4a**).

Collectively, our data suggest that H3 residues A1, R2 and K4 are the major determinants of H3 peptide binding, with smaller contributions from H3T3. Our recent study has also shown that H3Q5 is dispensable for binding as H3 Q5A peptide binds with similar affinity to WT H3^68^. These data suggest a finely tuned binding site with varying, methylation–dependent, contributions to H3 binding and H3K4 methyl discrimination. However, the contributions of these residues to binding may not be conserved between KDM5 family PHD1 domains. Mutations of H3K4 binding pocket residues in PHD1KDM5B, Y310 and L326 (Y294 and L310 in KDM5A) resulted in ∼2 and 4–fold lower affinity for unmodified H3 peptides was compared to WT^14^. In contrast, we observe much larger changes upon mutation of these residues, ∼15 to 60–fold change, in PHD1_KDM5A_, suggesting differential interactions of H3 with the PHD1 domains of KDM5A and KDM5B, despite their high sequence similarity. This is consistent with our structural work, which showed a difference in the binding mode of H3 to the PHD1 domain of KDM5A, and its binding to PHD1 of KDM5B^14^.

### Summary

Overall, our findings provide a structural understanding of how PHD1_KDM5A_ recognizes H3 tail residues and discriminates against methylated H3K4 peptides. Our findings demonstrate that the H3 peptide adopts a helical conformation that allows for extended interactions between the H3 ligand and PHD1 reader domain. In addition, the structures show unique interactions between N-terminal residues of PHD1 and H3K4, not previously observed in the PHD1 domains of KDM5 enzymes. Significantly reduced interactions of the N–terminal region of PHD1 with methylated H3K4 peptides further highlights the importance of this region in regulating the methylation selectivity in PHD1. We also identify the interactions that allow PHD1 to bind, with relatively high affinity, to methylated H3K4 peptides, a property that makes PHD1_KDM5A_, so far, unique when compared to related PHD domains. Given the allosteric role of this reader domain in KDM5A demethylase activity, our findings provide insights into how mutations in this domain would impact allosteric ligand binding and therefore modulate activity of KDM5A. The greater understanding of the structure provided here will facilitate development of PHD1–targeting small molecule modulators of KDM5A for use as allosteric regulators of demethylation.

## Acknowledgments

We would like to thank Nektaria Petronikolou and Fatima Ugur for their comments on the manuscript. In addition, we would like to thank Vicky Higman, Brian Smith, Benjamin Bardiaux and Ryan Tibble for their advice and NMR expertise.

