## Supplementary Figures 1-8 for "Recognition of histone H3 methylation states by the PHD1 domain of histone demethylase KDM5A"

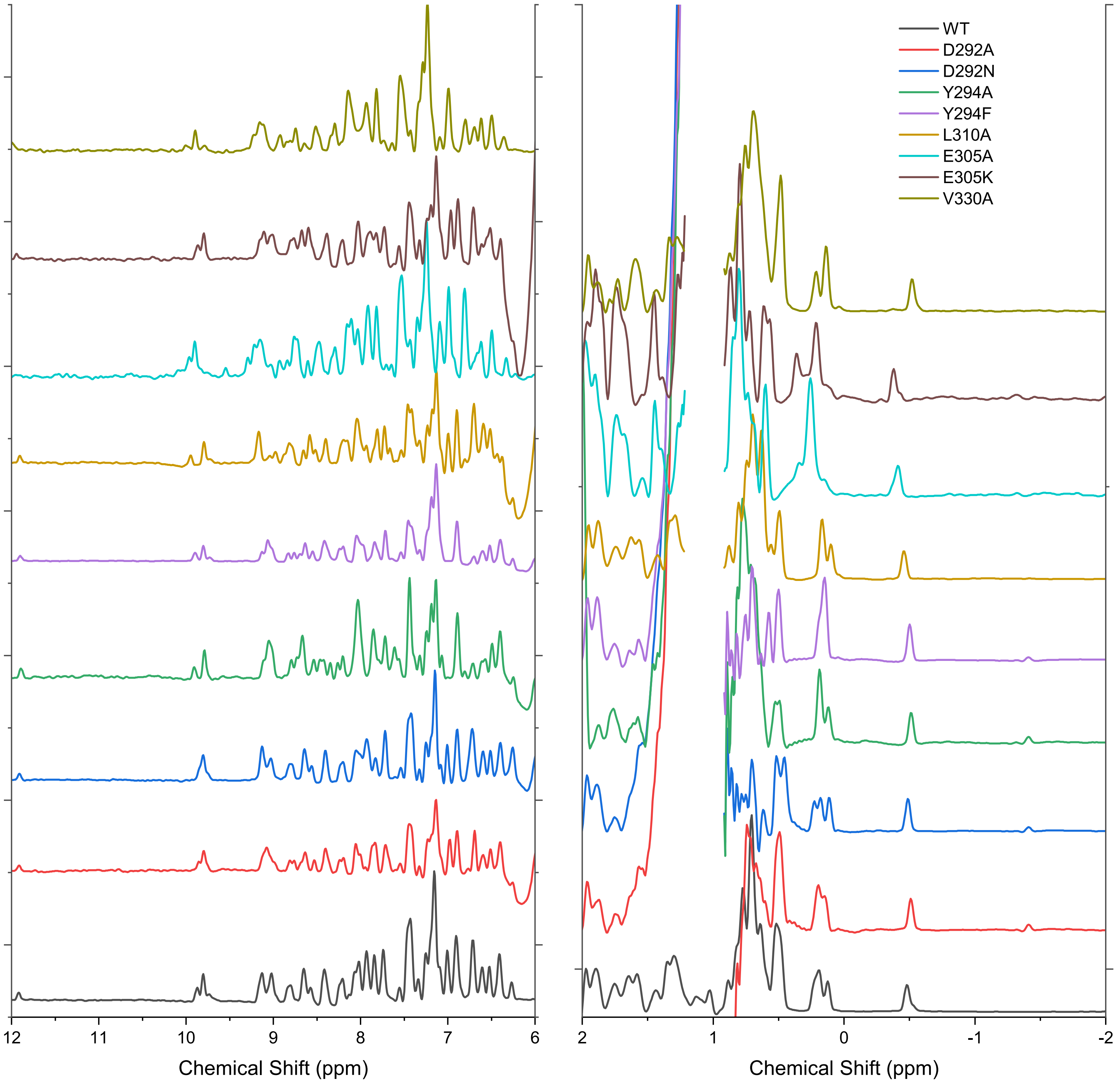


**Supplementary Figure 1.** 1H 1D NMR spectra of WT and mutant PHD1.

a

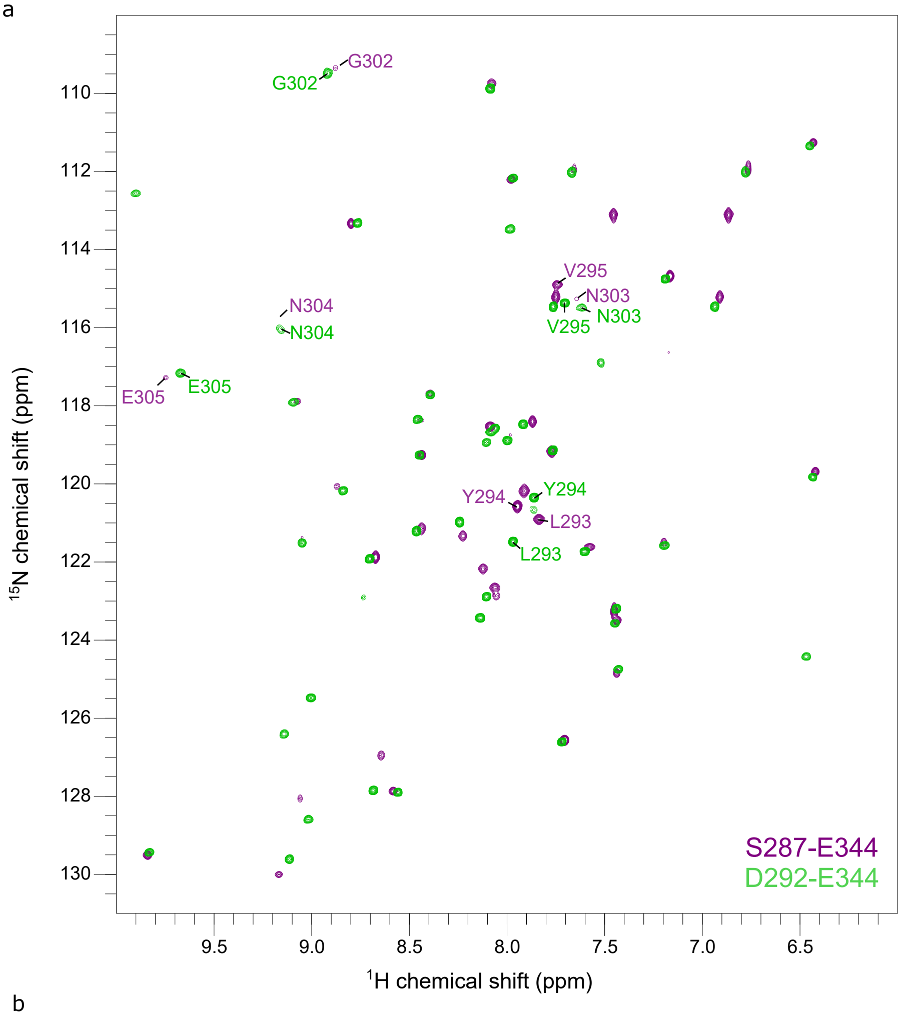


b

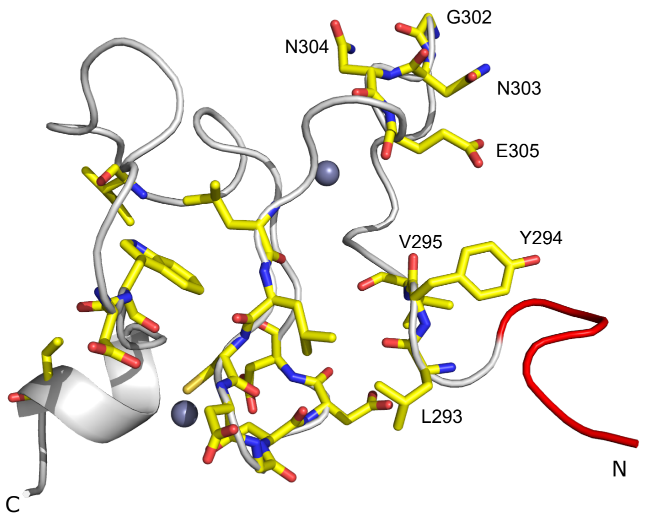


**Supplementary Figure 2.** a) Comparison of HSQC spectra of PHD1 S287-E344 (Purple) and D292-E344 (Green). b) Structure of apo PHD1 S287-E344 with residues that show a chemical shift difference between constructs shown in yellow. Red indicates residues not present in D292-E344. Specific N-terminal residues are highlighted.

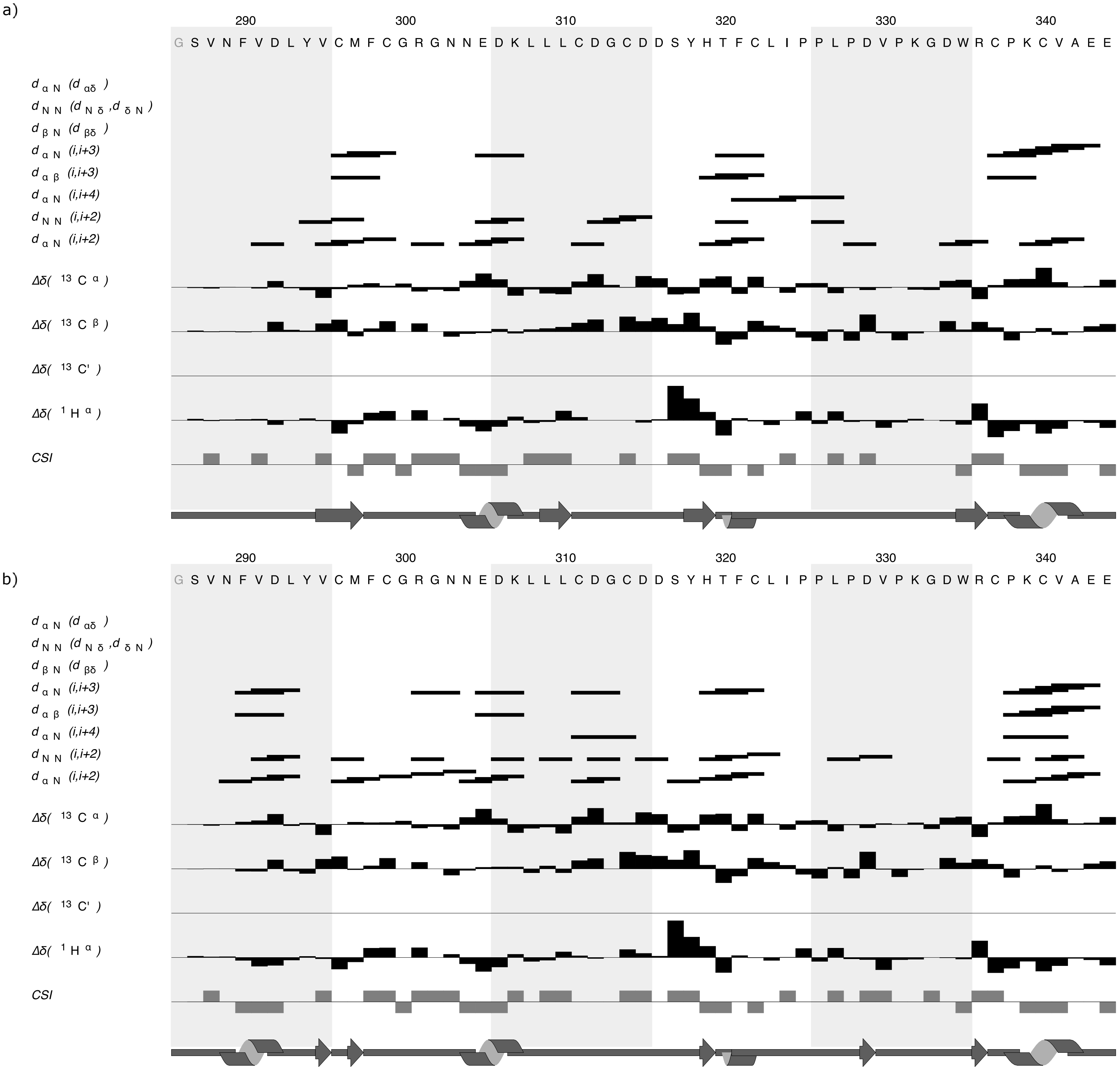


**Supplementary Figure 3.** Secondary structure prediction of a) Apo and b) H3 bound PHD1. Data were generated by CCPNMR analysis 2.4.2 secondary structure prediction tools.

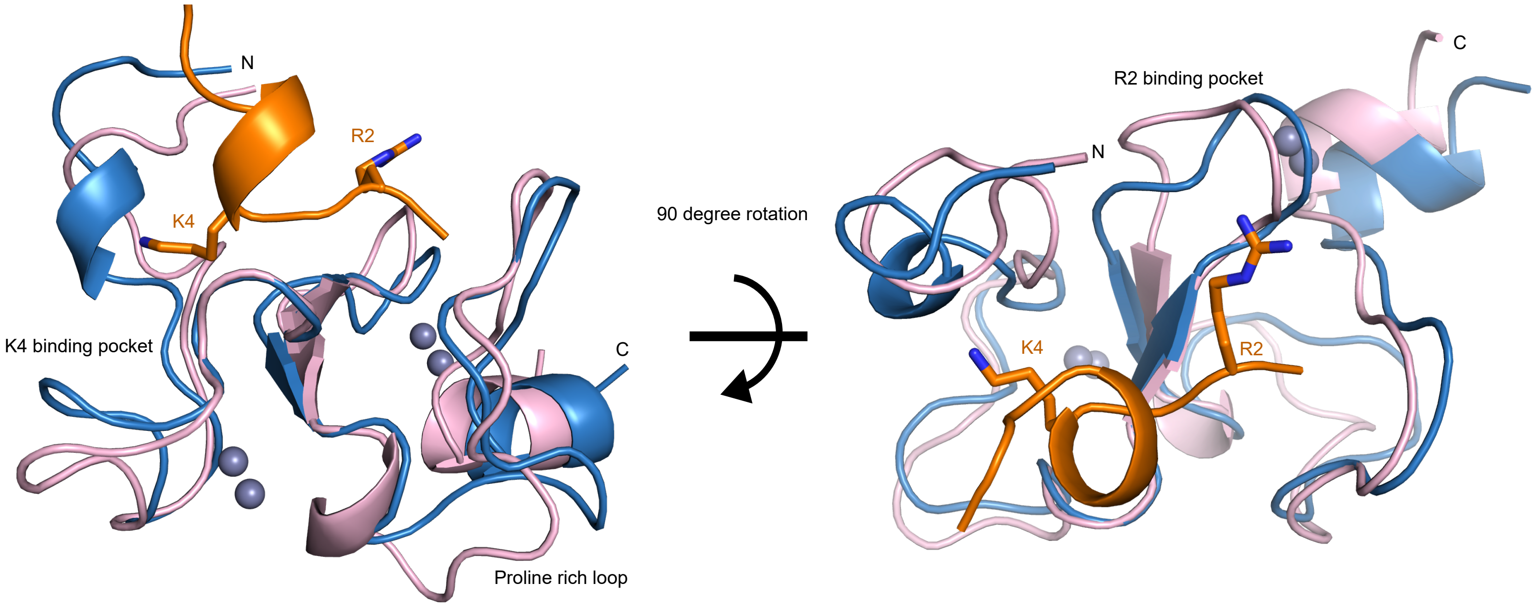


**Supplementary Figure 4.** Comparison between the apo (pink) and H3 bound structure of PHD1 (blue). H3 peptide is shown in orange. In the apo data set it was observed that for some residues, in addition to the major peak, a minor peak was present, which could suggest conformational flexibility. These minor peaks were absent in the H3 bound spectra.

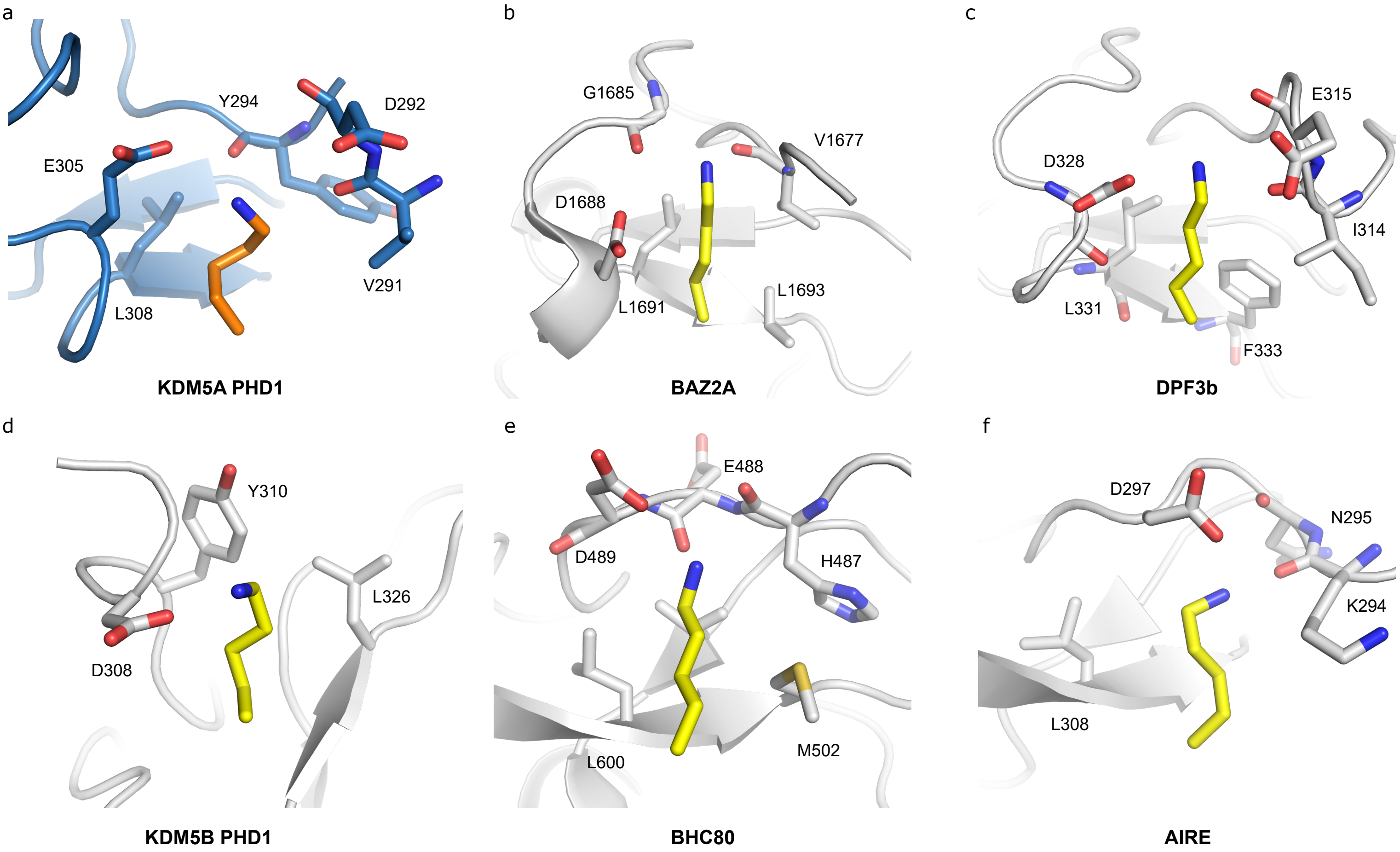


**Supplementary Figure 5. H3K4 binding pockets of different PHD domains.** a) PHD1 of KDM5A. b) PHD of BAZ2A (PDB: 5T8R). c) DPF3b (PDB: 5I3L). d) PHD1 of KDM5B (PDB: 2MNZ). e) BHC80 (PDB: 2PUY). f) AIRE (PDB: 2KFT). PHD1 of KDM5A is shown in blue and H3K4 is shown in orange. For all other structures the PHD or double PHD domain is shown in grey, H3K4 is shown in yellow.

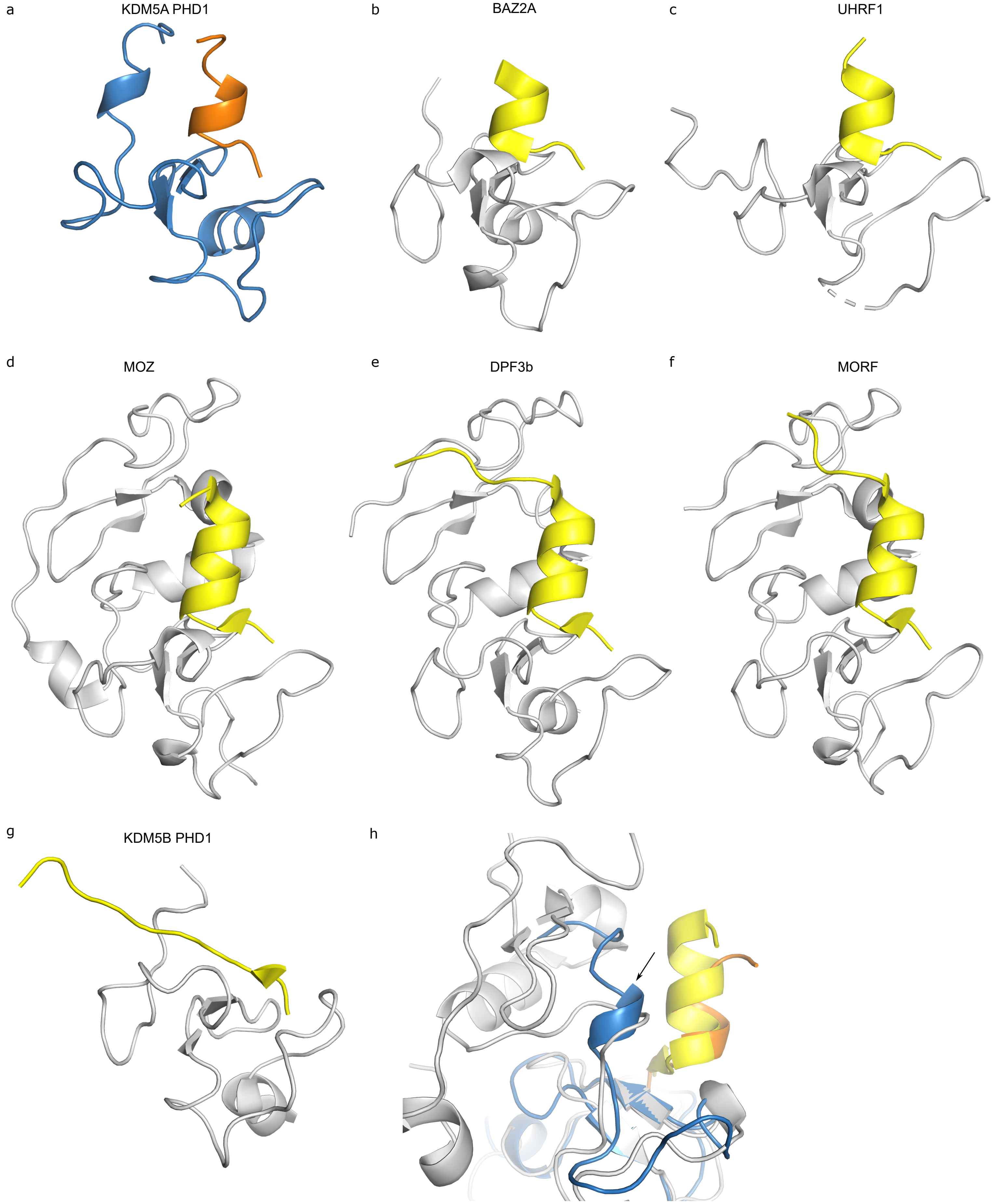


**Supplementary Figure 6.** a) PHD1 of KDM5A. b) PHD of BAZ2A (PDB: 5T8R). c) UHRF1 (PDB: 3ASK) d) MOZ (PDB: 4LK9). e) DPF3b (PDB: 5I3L). f) MORF (PDB: 5U2J). g) KDM5B PHD1 (PDB: 2MNZ) h) Overlaid structures of PHD1 of KDM5A and the double PHD finger of MOZ (PDB: 4LK9). N-terminal helical turn region (V291-L293) is indicated with an arrow. PHD1 of KDM5A is shown in blue and H3 is shown in orange. For all other structrues the PHD or double PHD domain is shown in grey, H3 is shown in yellow.

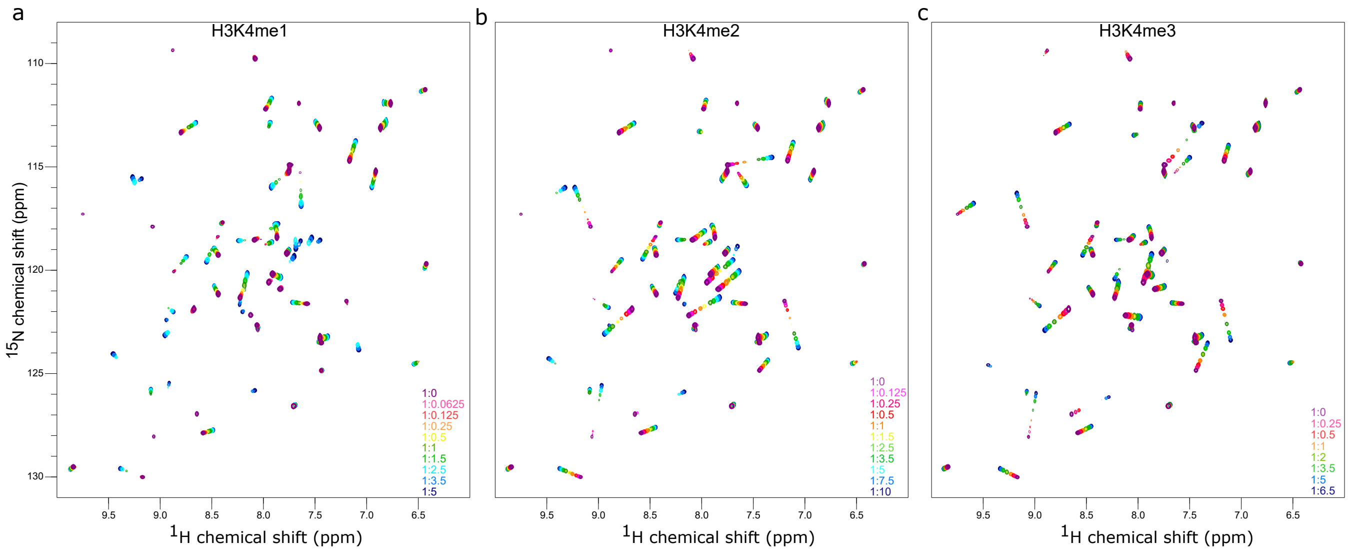


**Supplementary Figure 7.** HSQC spectra of H3K4me1/2/3 peptide titrations with 15N labeled PHD1 (S287-E344)

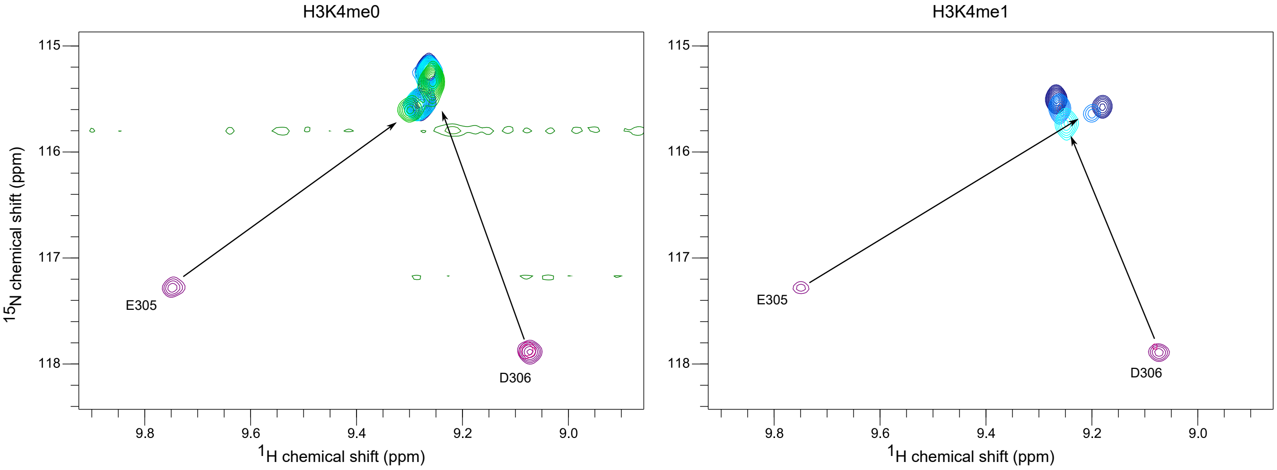


**Supplementary Figure 8.** Examples of slow exchanging peaks in HSQC titration experiments with unmodified H3 and H3K4me1 peptides
