## Supplementary Table 1 for "Recognition of histone H3 methylation states by the PHD1 domain of histone demethylase KDM5A"

**Supplementary Table 1.** NMR structure statistics

| | Apo PHD1 | PHD1 in complex with H3 10mer |
|---|---|---|
| **PDB ID** | 7KLO | 7KLR |
| **BMRB ID** | 30808 | 30809 |
| **Distance restraints** | | |
| Total NOE | 873 | 1265 |
| Intra-residue [i = j] | 348 | 563 |
| Sequential [\| i - j \| = 1] | 131 | 221 |
| Short range [2 ≤ \|i - j\| ≤ 3] | 58 | 103 |
| Medium range [4 ≤ \|i - j\| ≤ 5] | 53 | 55 |
| long range [\|i - j\| > 5] | 200 | 204 |
| Ambiguous | 83 | 119 |
| Hydrogen bonds (intra-/intermolecular) | 5 / 0 | 5 / 6 |
| Dihedral (psi/phi) | 45 / 45 | 48 / 48 |
| **Restraint statistics** | | |
| RMS of NOE violations (Å) | 0.319 ± 0.111 | 0.255 ± 0.094 |
| RMS of H bond violations (Å) | 0 | 0.665 ± 0.360 |
| RMS of dihedral violations (°) | 0.934 ± 0.087 | 0.717 ± 0.121 |
| **RMS from idealized covalent geometry** | | |
| Bonds (Å) | 0.00105 ± 0.00003 | 0.00124 ± 0.00002 |
| Angles (°) | 1.44 ± 0.07 | 1.32 ± 0.05 |
| Impropers (°) | 1.65 ± 0.08 | 1.32 ± 0.12 |
| **Average pairwise RMSD (Å)** | | |
| Heavy | 1.54 ± 0.31 | 1.6 ± 0.25 |
| Backbone | 1.12 ± 0.32 | 1.39 ± 0.27 |
| **Structure validation (PSVS)** | | |
| **Ramachandran plot** | | |
| Most favoured regions | 72.9 | 73.5 |
| Additionally allowed regions | 24.2 | 24.1 |
| Generously allowed regions | 2.9 | 0.2 |
| Disallowed regions | 0 | 2.2 |
| **Structure Quality Factors (raw/Z-scores)** | | |
| Verify3D | 0.03/-6.90 | 0.06/-6.42 |
| Prosall (-ve) | -002/-2.77 | 0.24/-1.70 |
| Procheck G-factor (phi-psi) | -0.95/-3.42 | -1.11/-4.05 |
| Procheck G-factor (all) | -0.71/-4.20 | -0.84/-4.97 |
| MolProbity clashscore | 9.69/-0.14 | 12.97/-0.70 |
